# Location-dependent proteomics of the aorta reveal an atherosclerotic disease gradient shaped by hemodynamics

**DOI:** 10.64898/2026.08.13.744640

**Authors:** Kathrine V. Jokumsen, Christina Christoffersen, Michael J. Davies, Luke F. Gamon

**Author notes:** Joint senior authors (Michael J. Davies, Luke F. Gamon). Corresponding authors *Email address:* (Luke F. Gamon), (Michael J. Davies).

## Abstract

**Background and aims:** Atherosclerotic plaques form preferentially at vascular sites exposed to disturbed blood flow, yet the protein changes underlying this site-specific plaque development remain unclear. Mouse models are widely used to study atherosclerosis but yield only limited amounts of tissue, previously restricting proteomic studies. However, recent advances in mass spectrometry now enable proteomic profiling of very small tissue samples. We aimed to utilise this to uncover site-specific protein changes in aortic regions prone or resistant to plaque formation.

**Methods:** Aortic arches from apolipoprotein E-deficient (ApoE^−/−^) mice fed a Western diet (WD) for 16 weeks were dissected into plaques from the major branches and inner curvature and visibly healthy regions. Proteins were extracted, enzymatically digested, and analysed by liquid chromatography-tandem mass spectrometry (LC-MS/MS).

**Results:** More than 4000 proteins were identified per sample despite their small size (< 1 mg tissue). Principal component analysis showed clustering by both disease status and anatomical location within the aortic arch, indicating distinct proteomes. Proteins known to drive atherosclerosis – including vascular cell adhesion molecule 1 (Vcam1), apolipoprotein B (Apob), lipoprotein lipase (Lpl), and galectin 3 (Lgals3) – were most abundant in advanced plaques and decreased progressively across anatomical regions, reaching their lowest levels in ‘healthy’ regions furthest from the plaques. Enrichment analysis highlighted pathways related to the extracellular matrix, immune system, hemostasis, and lipoprotein transport as central to disease progression.

**Conclusions:** This study demonstrates the feasibility of region-resolved proteomics in individual murine aortas and provide new molecular insights into the site-specific nature of atherosclerotic plaque development.

## 1. Introduction

Atherosclerosis is characterized by plaque development at specific sites of the vasculature. Hemodynamic factors largely dictate which vascular sites are susceptible or resistant to developing atherosclerosis [1–3]. Atherosclerosis tends to occur at sites of low shear stress and oscillatory or turbulent blood flow, which are found at branches and curvatures in the vasculature. Furthermore, developing plaques can themselves cause blood flow disturbances which may contribute to their growth over time [4]. These hemodynamic changes are perceived by endothelial cells lining the vasculature, which have been shown *in vitro* to respond by upregulating specific genes, such as Vcam1, NADPH oxidases, and nitric oxide synthases (NOS) [2, 5, 6]. However, there is a lack of *in vivo* data on the global protein changes occurring at specific vascular sites during plaque development. This knowledge gap is hindering the development of new therapeutics [7–9].

Proteomics is a powerful and unbiased analytical method that has been used to study atherosclerosis in mouse models (reviewed [7, 10]). However, earlier studies were constrained by limited protein coverage due to instrumental limitations and reported relatively few differentially regulated proteins, potentially due to pooling of samples from multiple animals [11–13]. A more recent study using a sequential extraction method detected a larger number of proteins and differentially expressed proteins, particularly extracellular matrix (ECM) proteins, which are challenging to analyse but are known to undergo substantial alterations during atherosclerosis progression [14]. Nevertheless, these studies analysed whole aortas, thereby lacking the spatial resolution needed to reveal site-specific proteomic changes associated with atherosclerotic plaque development. Spatial proteomics is an emerging field that enables regional mapping of tissue heterogeneity and may help uncover these spatial disease patterns that are obscured in bulk tissue analyses [15, 16].

In this study, we mapped the site-specific proteome of the aortic arch from ApoE^−/−^ mice dissected into regions prone or resistant to plaque formation. An efficient, single-step protein extraction procedure was used in combination with state-of-the-art LC-MS/MS-based proteomics. Major protein changes were detected between plaques and ‘healthy’ aortic areas, but also location-dependent changes between plaques.

## 2. Materials and methods

### 2.1. Animal experiments

The animal study was approved by the Danish Animal Experiments Inspectorate (approval number: 2021-15-0201-00818). 8-weeks-old male ApoE^−/−^ mice (B6.129P2-*Apoe*^*tm1Unc*^ N11) purchased from Taconic received a high fat Western diet (D12079Bi, Research Diets) and water *ad libitum* for 16 weeks (*n* = 5 mice). At sacrifice, animals were anesthetized with Zoletil, bled out through the retro-orbital venous sinus, and perfused with ice-cold saline through the left ventricle. The aortic arch, including the thoracic descending aorta, was isolated after removal of periaortic adipose tissue.

### 2.2. Dissection of the aortic arch

*En face* preparation of the aortic arch was performed to expose the intimal surface. Plaques were visualised under a microscope as thickened and calcified vessel wall and dissected from the three major branches and inner curvature of the aortic arch. Regions of visibly healthy artery wall were dissected as well from the aortic arch and descending aorta.

### 2.3. Sample preparation for LC-MS/MS-based proteomic analysis

Proteins were extracted from the aortic samples using the Sample Preparation by Easy Extraction and Digestion (SPEED) protocol [17], reduced and alkylated, digested with LysC and trypsin, and cleaned up by stage tipping [18]. Samples were the separated by LC and analysed using a Bruker timsTOF Pro mass spectrometer operated in a data-independent acquisition with parallel accumulation-serial fragmentation (DIA-PASEF) mode [19]. For further details see the Supplementary Data.

### 2.4. Data analysis

Data were searched against the UniProt protein database, including common contaminants, using DIA-NN (version 1.9.2) [20] in library-free mode (see Suppl. Data). DIA-Analyst (version 0.10.3, https://analyst-suites.org/apps/dia-analyst) was used for standardised downstream statistical analysis of the protein-level data. Proteins were considered significantly regulated if the Benjamini-Hochberg (BH) [21] adjusted p-value was < 0.05 and log_2_ fold change > 1. Further data analysis and visualisation was performed using R (version 4.5.2). Intensity based absolute quantification (iBAQ) values were calculated as described previously [22].

## 3. Results

### 3.1. Proteomic profiling of plaques and ‘healthy’ areas of the aortic arch from ApoE^−/−^ mice

To study the site-specific nature of atherosclerotic plaque development by proteomics, plaques and ‘healthy’ areas of the aortic arch from ApoE^−/−^ mice fed a high-fat WD were dissected (Fig. 1A). Plaques were consistently detected in the three principal branches of the aortic arch (the brachiocephalic, left common carotid, left subclavian artery; designated P1, P2 and P3) and the inner curvature (P4), with the first branch, P1, containing the largest plaque. This is consistent with previously identified sites of plaque development [23]. ‘Healthy’ areas of artery wall were dissected from the ascending aorta and arch (A1), as well as descending aorta (A2 and A3). These samples were subjected to LC-MS/MS-based proteomic analysis as outlined in Fig. 1B.

**Fig. 1.**
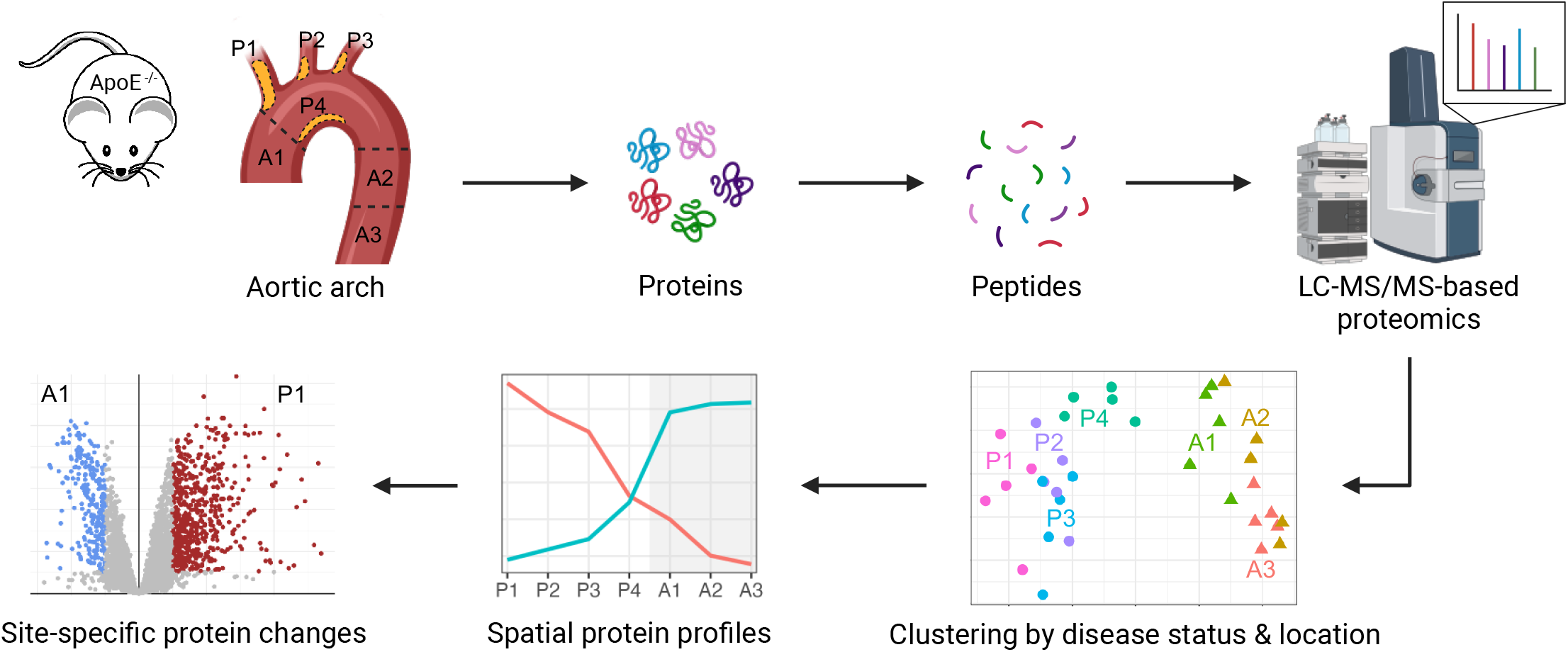
Mapping site-specific aorta proteomes. **(A)** The aortic arch was isolated from atherosclerosis-prone ApoE^−/−^ mice fed a high-fat Western diet (WD) for 16 weeks (*n* = 5). En face preparation was done to expose the intimal vessel surface with the atherosclerotic plaques (white). Plaques were dissected (as outlined in red) from the 3 principal branches, the brachiocephalic artery (P1), the left common carotid artery (P2), and the left subclavian artery (P3), as well as the inner curvature of the aortic arch (P4). Areas of unaffected ‘healthy’ artery wall were also dissected (outlined in green) from the ascending aorta and arch (A1), along with upper (A2) and lower (A3) parts of the descending thoracic aorta. **(B)** Proteins were extracted from these aortic samples using the SPEED protocol, reduced and alkylated, digested with LysC / trypsin, cleaned up by stage tipping, and analysed by liquid chromatography-tandem mass spectrometry (LC-MS/MS). **(C)** The number of proteins identified in each of the 7 areas of the aortic arch. Note colour coding according to aortic area. **(D)** Sample correlation heatmap with hierarchical clustering. **(E)** Principal component analysis (PCA) revealed clear separation of plaque and ‘healthy’ aorta tissue and additional separation by location within each of the two clusters. **(B)** was created with BioRender.com/w31f4c9.

A total of 4856 proteins were identified and quantified across all samples. Slightly more proteins were identified from the plaques (P1-P4; 4360) than the ‘healthy’ aorta samples (A1-A3; 4133 proteins) (Fig. 1C). A large overlap in proteomes was observed between the 7 different areas of the aortic arch, with 4404 commonly identified proteins (Suppl. Fig. 1), suggesting that it is predominantly the differential expression of common proteins rather than the presence or absence of specific proteins that underlie site-specific plaque development. This was further supported by sample correlation in protein expression patterns (Fig. 1D). Pearson correlation coefficients were consistently high (> 0.84) among samples of the same disease status, but lower (0.62 – 0.96) between plaques and ‘healthy’ aorta samples, consistent with biological differences in protein expression profiles. A gradient pattern was observed across plaque locations where P1 showed the lowest correlation with ‘healthy’ aorta samples, P2 and P3 intermediate, and P4 the highest. This indicates a progressive difference in protein expression profiles, with P4 most closely resembling the ‘healthy’ aorta. Similarly, A1 showed a higher correlation with the plaques than A2 and A3.

A similar pattern emerged from principal component analysis (PCA) (Fig. 1E). Disease status was the main driver of sample separation, as the plaque and ‘healthy’ aorta samples formed two distinct clusters. However, a secondary separation by aortic location was also apparent within each cluster, as P1 and P4 forming distinct subclusters from P2 and P3, and A1 separated clearly from A2 and A3. Most of the PCA separation was driven by principal component 1 (PC1) with the top 50 protein loadings contributing to PC1 reported in Suppl. Table 1. These included several lipoprotein-associated proteins (Fig. 2A), proteins involved in immune cell adhesion and activation (Fig 2B), macrophage proteins (Fig 2C), protease inhibitors (Fig. 2D), complement proteins (Fig 2E), fibrinogen (Fig. 2F), ECM proteins (Fig. 2G), a myosin protein (Fig. 2H), and multiple immunoglobulins (Suppl. Fig. 2). The protein expression profiles of these species across the anatomical areas were largely similar, displaying a gradient with highest expression in P1–P3, a gradual decrease through P4 and A1, and low levels in A2 and A3. Several of these proteins are established drivers of atherosclerosis, including Apob [24], Vcam1 [25], CD5 antigen-like (Cd5l) [26], and Lgals3 [27], or abundant plasma proteins known to be concentrated in atherosclerotic intima, such as fibrinogen, complement proteins, and immunoglobulins [28]. The only protein of the top 50 PC1 loadings that displayed an opposite expression profile was myosin light chain 6B (Myl6b), which had its lowest expression in P1 and highest in A3. These findings indicate that there is a significant gradient in atherosclerosis-related proteins across the aortic arch, with P1 having the most advanced plaque, P4 showing features of an early plaque that is most similar to ‘healthy’ aorta, and A1 being the most atherosclerosis-prone area of ‘healthy’ aorta (i.e. atherosclerosis severity: P1 > P2 > P3 > P4 > A1 > A2 ≥ A3).

**Fig. 2.**
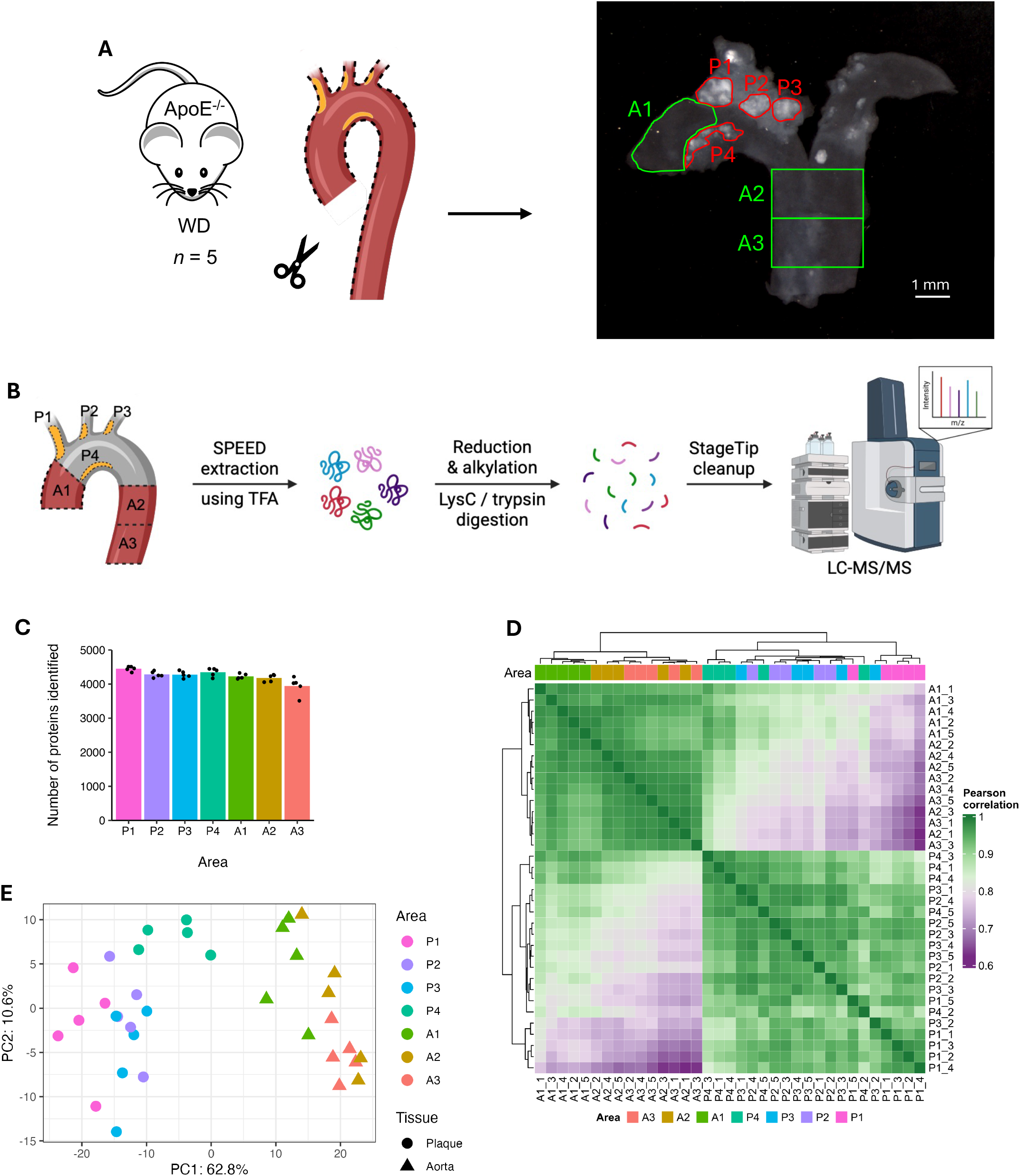
Protein expression profiles across aortic areas of the top 50 protein loadings of principal component 1 (PC1; see Fig. 1E). These include: **(A)** lipoprotein-associated proteins apolipoprotein B-100 (Apob), apolipoprotein C-I (Apoc1), lipoprotein lipase (Lpl), serum paraoxonase/arylesterase 1 (Pon1); **(B)** proteins involved in immune cell activation and adhesion like CD63 antigen (Cd63), secreted frizzled-related protein 3 (Frzb), integrin beta-2 (Itgb2), plastin-2 (Lcp1), P2X purinoceptor 4 (P2rx4), vascular cell adhesion molecule 1 (Vcam1); **(C)** macrophage proteins CD5 antigen-like (Cd5l), galectin-3 (Lgals3), neutral cholesterol ester hydrolase 1 (Nceh1); **(D)** protease inhibitors inter-alpha-trypsin inhibitor heavy chain 1, 2 & 4 (Itih1, Itih2 & Itih4), murinoglobulin-1 (Mug1), pregnancy zone protein (Pzp); **(E)** complement proteins C1q subunit A & B (C1qa & C1qb), C3 (C3), C4b-binding protein (C4bpa), mannan-binding lectin serine protease 2 (Masp2), mannose-binding protein A (Mbl1), **(F)** fibrinogen alpha, beta & gamma chain (Fga, Fgb & Fgg); **(G)** extracellular matrix (ECM) proteins collagen XII alpha-1 chain (Col12a1), fibromodulin (Fmod), fibronectin (Fn1), matrix Gla protein (Mgp), thrombospondin-1 (Thbs1), tenascin (Tnc); and **(H)** myosin light chain 6B (Myl6b). Data are presented as mean log2 intensity ± SEM from *n* = 5 mice.

### 3.2. Site-specific protein changes in the aortic arch

Subsequently, differential protein expression was investigated between different areas of the aortic arch susceptible or resistant to atherosclerosis. Firstly, differential expression analysis was carried out between the most advanced plaque P1 and the nearby ‘healthy’ area A1. 856 differentially expressed proteins were identified between P1 and A1, with the majority of these being upregulated in P1, consistent with a high degree of difference between plaque-containing and visibly healthy artery wall (volcano plot in Fig. 3A; a full list of proteins is provided in Suppl. Table 2). The likely cellular origin of these protein changes was examined by enrichment analysis using cell type signature gene sets from single-cell sequencing studies [29–31]. Multiple different immune cells were enriched in P1, including macrophages, monocytes, Natural Killer (NK) cells, T cells and B cells (rug plots in lower part of Fig. 3A). Fibroblasts were also enriched in P1, whereas muscle cells were enriched in A1.

**Fig. 3.**
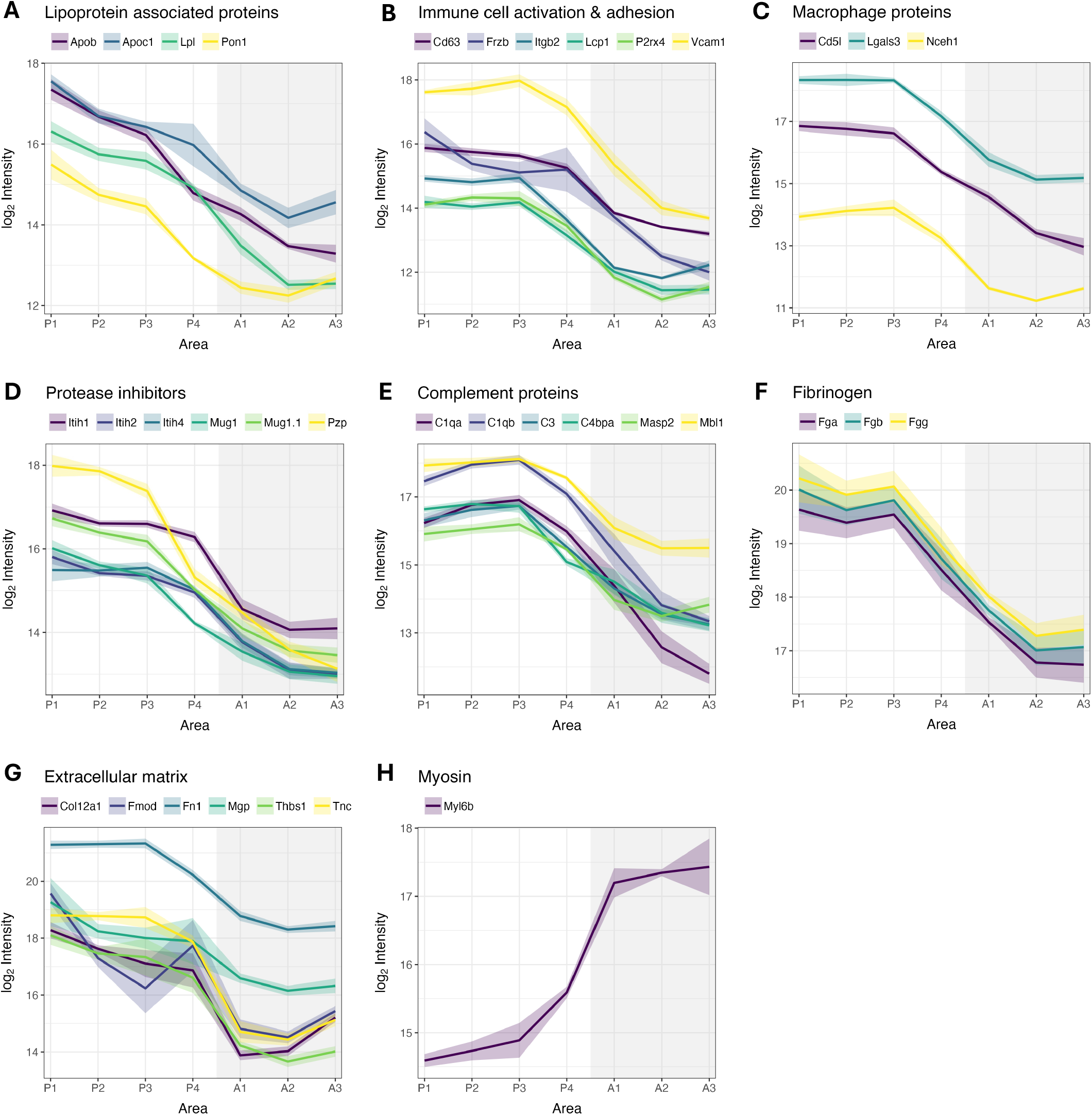
Location-specific aorta protein signatures. **(A)** Volcano plot with rug plot showing result of differential expression analysis followed by gene set enrichment analysis of cell type signatures comparing plaque P1 to nearby ‘healthy’ aorta area A1. Proteins were considered significant if adjusted *p* value < 0.05 and log2 fold change > 1. Blue: upregulated in A1, red: upregulated in P1, grey: non-significant. **(B – F)** Volcano plots comparing P1 to the five other ‘healthy’ and plaque regions of the aortic arch. Proteins differentially regulated across all six comparisons are highlighted in panel **(F). (G)** Distribution of neutrophil proteins across all aortic areas (*n* = 5 mice): neutrophil gelatinase-associated lipocalin (Lcn2), plastin-2 (Lcp1), myeloperoxidase (Mpo), neutrophilic granule protein (Ngp), protein S100-A8 (S100a8), protein S100-A9 (S100a9). **(H)** Ranked protein abundance in P1 with top 50 most abundant neutrophil proteins highlighted in green.

P1 was also compared to the five other areas of the aortic arch (Fig 3B-F). The number of differentially regulated proteins was greatest when P1 was compared to ‘healthy’ areas (Suppl. Fig. 3), particularly A2 and A3 of the descending aorta, with 613 proteins overlapping in the comparisons between P1 and A1, A2 or A3. However, differentially expressed proteins were also identified between P1 and plaques from other locations in the aortic arch, with the greatest number (311) between regions P1 and P4 (Fig. 3D, Suppl. Fig. 3). This is consistent with P4 being the least advanced plaque (of P1-P4), closest to ‘healthy’ aorta. 265 of the differentially expressed proteins between P1 and P4 overlapped with those between P1 and A1, consistent with the close proximity of A1 and P4. The comparison of P1 with P2 (Fig. 3F) and P3 (Fig. 3E) (i.e. the two other branches of the aortic arch) revealed 29 and 72 differentially expressed proteins, respectively (Suppl. Fig. 3). This indicates location-dependent differences in protein expression between plaque areas. However, most of the plaque proteome remained unchanged across locations, with around 4800 proteins showing no significant difference in expression between plaque locations, suggesting a high degree of plaque proteome conservation.

Twelve proteins (highlighted in Fig. 3F) were consistently differentially regulated in P1 when compared to the six other aortic areas. These include the neutrophil-specific proteins neutrophilic granule protein (Ngp) and protein S100-A9 (S100a9). Only a small neutrophil gene set was identified in the cell type enrichment analysis (Suppl. Table 3), probably reflecting the scarcity of neutrophil gene sets in the collection used. Closer inspection of the common neutrophil proteins revealed that these were not only most abundant in P1, but often only consistently detected in the plaques (Fig. 3G). Ranking of all proteins by abundance in P1 and highlighting the most abundant neutrophil proteins [32] showed that these were in the moderate to high abundance range (Fig. 3H). These data are consistent with neutrophils being present in murine atherosclerotic plaques.

Pathways associated with these site-specific protein changes were examined by gene set enrichment analysis using the fold changes from the comparisons of P1 to the six other aortic areas. This revealed a considerable overlap with several pathways related to ECM organization, hemostasis, the innate immune system, and insulin-like growth factor (IGF) being enriched in P1 in all or most comparisons (Suppl. Fig. 4). It included specific pathways such as ECM degradation, integrin cell surface interactions (with ECM), collagen formation, fibrin clot formation, cell surface interactions at the vascular wall (leukocyte extravasation), toll-like receptor cascades, and neutrophil degranulation. Other pathways were only enriched in P1 when compared to the ‘healthy’ areas (A1-3) and P4, suggesting that these are key pathways in plaque development and progression. This includes pathways related to the immune system (complement cascade, ROS / RNS production in phagocytes), plasma lipoprotein assembly, remodelling and clearance, iron uptake and transport, protein translation, RNA metabolism (nonsense-mediated decay and rRNA processing), and cellular response to stress (amino acids regulate mTORC1). Pathways that were negatively enriched, or downregulated, in P1 revolved around muscle contraction, signalling by Rho GTPases, metabolism (particularly aerobic respiration and respiratory electron transport), and cell-cell communication (including cell junction organization and cell-ECM interactions). These data are consistent with phenotypic switching of smooth muscle cells from a contractile to a synthetic type, a metabolic shift from aerobic respiration to anaerobic glycolysis, and loss of cell-cell contact during endothelial-mesenchymal transition.

### 3.3. Extracellular matrix proteins in atherosclerotic plaques and healthy aorta

The enrichment of several ECM-related pathways was of interest, as the ECM has been particularly difficult to study previously but is known to play an important role in atherosclerosis [33]. Further analysis showed that core ECM proteins (as defined by [34]) were fewer in number but more abundant than cellular and plasma proteins (163 core ECM proteins versus 4470 cellular and plasma proteins; Suppl. Fig. 5), as indicated by their iBAQ values (Fig. 4A). Core ECM proteins were the most abundant species in all samples analysed (Suppl. Fig. 6), indicating that core ECM proteins are an important constituent of all regions of the aorta irrespective of the presence or absence of plaque. ECM-associated proteins were more abundant than cellular and plasma proteins in plaques, but not ‘healthy’ aorta. However, when considering all aortic areas separately, ECM-associated proteins were more abundant in all areas except A3. This suggests that proteins which interact with or modify the ECM are important during atherogenesis.

**Fig. 4.**
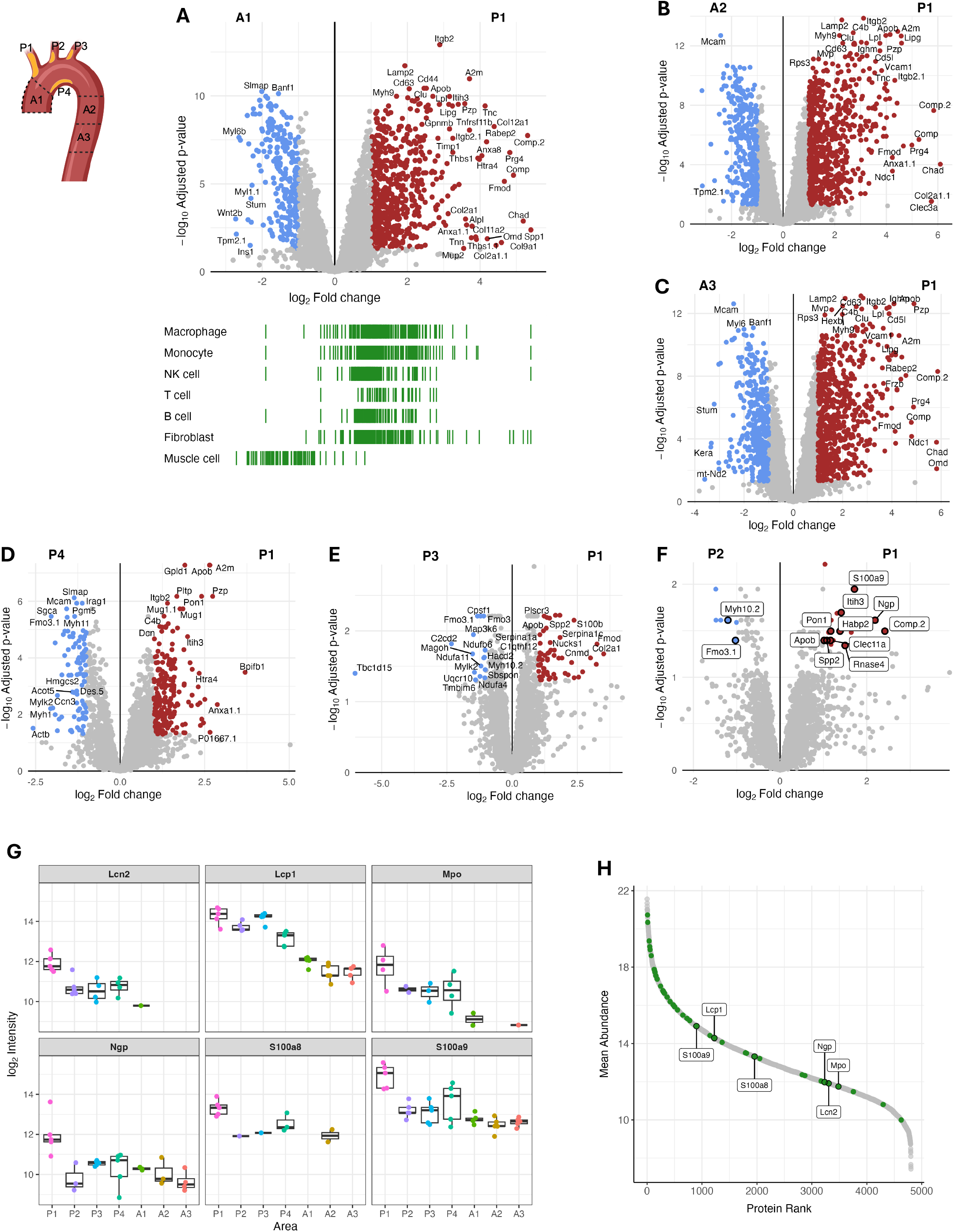
Extracellular matrix (ECM) proteins in plaque and ‘healthy’ aorta. **(A)** Estimated absolute abundance of core ECM, ECM-associated, and cellular and plasma proteins in all ‘healthy’ aorta and plaque samples, respectively. Violin plots show the distribution of iBAQ values, with crossbars indicating the mean abundance. Statistical significance was determined using a BH-adjusted *t* test. ^*^ *p* < 0.05, ^**^ *p* < 0.01, ^***^ *p* < 0.001. **(B)** Volcano plot comparing plaque P1 to ‘healthy’ aortic area A1 with differentially regulated ECM proteins annotated. Proteins were considered significant if adjusted p value < 0.05 and log2 fold change > 1. Pink: core ECM proteins; green: ECM-associated proteins; dark grey: cellular and plasma proteins; light grey: non-significant proteins. **(C)** Mean normalized protein expression across aortic areas of the 6 ECM protein categories. Individual proteins grouped based on similar protein expression profiles. The groups showed distinct patterns of protein expression which were comparable between the 6 ECM categories, as they either decreased (red) or increased (blue) in expression going from P1 to A3. The collagens could be divided into three groups, with the third having a dual peak expression in P1 and P4.

**Fig. 5.**
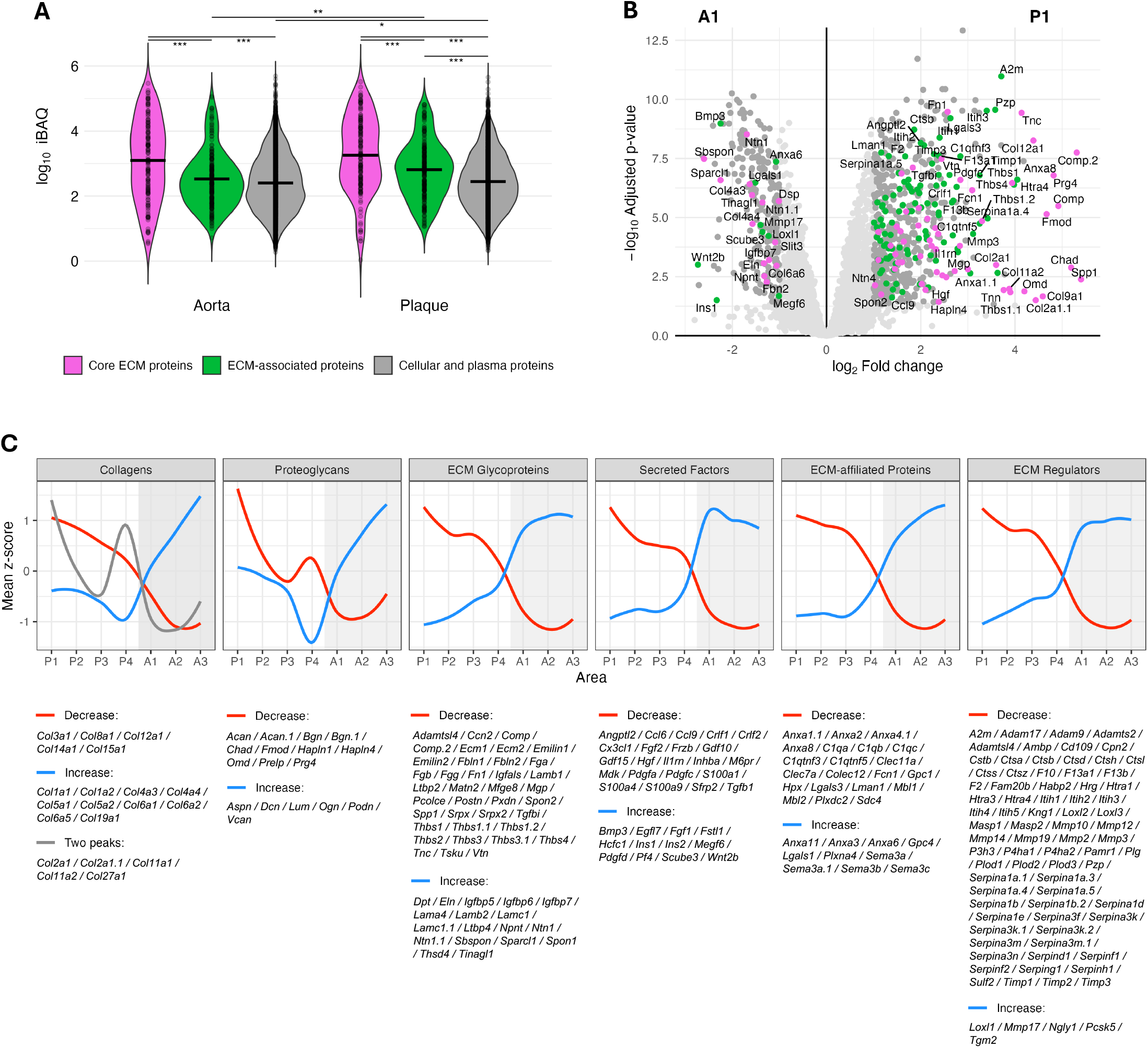

Of the differentially regulated proteins identified between P1 and A1, 68 were core ECM proteins and 109 were ECM-associated (Fig. 4B). Notably, 46% of the most upregulated proteins in P1 (log_2_ fold change > 2) were ECM proteins. This evidence for site-specific differences in ECM protein expression prompted further analysis of the protein expression profiles across the 7 aortic areas. Core ECM proteins were categorized as either collagens, ECM glycoproteins or proteoglycans, while ECM-associated proteins were divided into secreted factors, ECM-affiliated proteins and ECM regulators. No distinct trends in protein expression across aortic areas was observed for these 6 ECM protein categories, apart from the ECM regulators (Suppl. Fig. 7). However, further analysis of individual proteins in each category revealed that these could be divided into two (or three) groups based on similar protein expression profiles. These groups showed distinct patterns in protein expression across the aortic areas, which were comparable between the 6 ECM categories (Fig. 4C). The first group gradually decreased in protein expression from P1 to A2, with the most pronounced decrease occurring between P4 and A1. Some of the proteins in this group have been previously implicated in atherosclerosis, including collagen VIII alpha-1 chain (Col8a1) [35], fibronectin (Fn1) [36], and peroxidasin (Pxdn) [14]. The majority of the ECM regulators followed this expression profile, and included multiple proteases (e.g. cathepsins, matrix metalloproteinases, serine proteases), protease inhibitors (e.g. serpins, metalloproteinase inhibitors, inter-alpha-trypsin inhibitor), and collagen-modifying enzymes (Fig. 4C). The second group gradually increased in protein expression from P1 to A3, with the greatest increase occurring between P4 and A1. This group included elastin, collagen types I, V and VI, and laminins which are major ECM components of normal blood vessels (Fig. 4C) [37].

Interestingly, the proteoglycans showed more complex protein expression profiles than the other ECM categories with a specific peak or trough at P4. The collagens divided into a third group which decreased in protein expression from P1 to P3, then increased at P4, before decreasing again in the ‘healthy’ aortic areas (Fig. 4C). This dual peak of high expression in P1 (most advanced plaque) and P4 (least advanced plaque), indicates that this group of collagens and proteoglycans might be important at several different stages of plaque development.

## 4. Discussion

Atherosclerotic plaques develop at specific vascular sites exposed to blood flow disturbances, but the mechanisms underlying this spatial disease pattern remain incompletely understood. Previous studies have identified a limited number of proteins that are changed in artery wall regions exposed to low shear stress and oscillatory or turbulent flow and have associated these protein changes with plaque development (reviewed [1–3]). However, untargeted studies examining global protein changes across different arterial sites that are susceptible or resistant to plaque development remain limited (reviewed [7, 10]). Furthermore, most previous murine proteomic studies have analysed whole aortas, resulting in a loss of spatial resolution and masking regional differences in protein expression [11, 12, 14].

Here, we employed state-of-the-art LC-MS/MS-based proteomic methods to explore the site-specific nature of atherosclerotic plaque development. We quantified 4856 proteins across different regions of the aortic arch from ApoE^−/−^ mice with or without visibly plaque formation, of which 1440 proteins were differentially expressed between regions. Together, these results show that our approach of microdissection combined with single-step SPEED protein extraction provides comprehensive and spatially resolved proteome coverage of the murine aorta, with particularly good coverage of ECM proteins.

PCA and sample correlation revealed pronounced proteome differences between atherosclerotic plaques and visibly healthy aortic areas, as expected (Fig. 1D-E). However, anatomical location within the aortic arch also affected the proteomes substantially. Plaques from the four different locations in the aortic arch differed in protein composition, with P4 (inner curvature) showing the highest similarity to ‘healthy’ aorta, P2 and P3 intermediate (2^nd^ and 3^rd^ branch), and P1 (1^st^ branch) the lowest. This supported the hypothesis that P1 is the most advanced and P4 is the least advanced atherosclerotic plaque. For the visibly healthy aortic regions, the proteome of A1 (ascending aortic and arch) was more similar to the plaques than A2 and A3 (descending aorta), indicating that A1 is a more atherosclerosis-prone region despite the absence of visible plaque. The proteins contributing most strongly to these proteome differences exhibited remarkably similar expression profiles, forming a gradient with highest expression in P1 and lowest in A3 (Fig. 2A-G; Suppl. Fig. 2). Several of these proteins are well-known drivers of atherosclerosis, including Apob [24], Vcam1 [25], Cd5l [26], and Lgals3 [27], indicative of a molecular disease gradient across the aortic arch following the sequence P1 > P2 > P3 > P4 > A1 > A2 ≥ A3. In addition, these top protein loadings included several lipoprotein-associated proteins, immunoglobulins, complement proteins, fibrinogen, and protease inhibitors, all of which are abundant plasma proteins [28]. This suggests that a gradual increase in permeability and retention of plasma proteins in the artery wall is associated with the site-specific plaque development in the aortic arch. This interpretation aligns with previous studies showing a continuous decrease in aortic permeability to LDL with increasing distance from the heart [38]. These findings, together with the established upregulation of Vcam1 in response to low shear stress and oscillatory blood flow, support a role for local hemodynamic forces in shaping site-specific plaque development and associated proteome changes [2, 5, 25].

The protein expression changes between plaque P1 and ‘healthy’ aortic areas were overwhelmingly upregulations in P1, with only few proteins being downregulated (Fig. 3A-C). This asymmetry likely reflects the development of the atherosclerotic plaque in the vessel wall, which involves an accumulation of lipoprotein particles, ECM-producing smooth muscle cells, and infiltrating immune cells. The latter was supported by the cell type enrichment analysis, which showed that multiple immune cell types were enriched in P1, including macrophages, monocytes, NK cells, T cells, and B cells (Fig. 3A). This is consistent with previous studies demonstrating that all of these immune cell types accumulate in atherosclerotic plaques [39]. These observations were further supported by the marked upregulation of proteins involved in leukocyte adhesion and extravasation, including integrin beta-2 (Itgb2) and intercellular adhesion molecule 1 (Icam1) (Fig. 3A-C, Suppl. Table 2) [40].

Although neutrophils were only weakly identified in the cell type enrichment analysis (Suppl. Table 3), this likely reflects the scarcity of neutrophil gene sets in available single-cell transcriptomic reference datasets. Such underrepresentation is unsurprising given that neutrophils are technically challenging to characterize using transcriptomics [41]. In contrast, neutrophil-associated proteins were upregulated in P1 (Fig. 3G), exhibited moderate to high abundance (Fig. 3H), and neutrophil degranulation was among the pathways enriched in P1 (Suppl. Fig. 4). Collectively, these findings suggest that neutrophils are an important cellular component of advanced murine atherosclerotic plaques and may contribute to disease progression. This interpretation is consistent with experimental data from human plaques [16, 42–44] and epidemiological studies [45], despite the lower levels of neutrophils in mice (10-25% of total leukocytes) than in humans (50-70%) [46]. Notably, the upregulation of neutrophil proteins S100a8 and S100a9 in P1 is consistent with a previous proteomics study that identified these proteins as key features of unstable plaques in a tandem stenosis mouse model [13]. This raises the possibility that plaques in the brachiocephalic artery of WD-fed ApoE^−/−^ mice, despite their distinct morphology, share molecular characteristics with unstable plaques and may therefore be more similar than previously appreciated [47, 48].

A particular strength of this study is the extensive coverage of ECM proteins achieved by through the use of the single-step SPEED extraction protocol [17]. Owing to technical challenges in the extraction and detection of ECM proteins, the ECM proteome has historically been underrepresented in proteomic datasets, particularly in mouse models, potentially leading to an underestimation of its importance (though see [42, 49] for human data). The present dataset therefore provides extensive spatially resolved coverage of ECM proteins across plaque-prone and -resistant areas of the aortic arch. The importance of ECM proteins was highlighted by the observation that they represented the most abundant protein class across all aortic regions, irrespective of plaque status (Fig. 4A; Suppl. Fig. 6). ECM proteins represented almost half (46%) of the proteins most strongly upregulated in P1 relative to A1 (log_2_ fold change > 2; Fig. 4B), despite accounting for only ~8% of the total quantified proteins (Suppl. Fig. 5). The ECM proteins followed one of two reciprocal expression profiles across the aortic arch, either decreasing or increasing in expression from P1 to A3 (Fig. 4C). The first group of proteins were most abundant in plaques and gradually decreased across locations. This group included Col8a1, Fn1, and Pxdn, which have previously been implicated in atherosclerosis [14, 35, 36], suggesting a potential association with plaque development. Multiple proteases and protease inhibitors also followed this expression profile, being most abundant in P1, which suggests enhanced ECM degradation and remodelling. This interpretation is consistent with studies of unstable human plaques, which have detected large numbers of proteases and protein fragments resulting from their activity [42, 50]. Collectively, these spatially resolved proteomic data indicate that plaques in ApoE^−/−^ mice, particularly those within the brachiocephalic artery, share molecular similarities to unstable human lesions.

An important consideration when interpreting these findings is the substantial overlap in protein identifications across the 7 aortic areas (Suppl. Fig. 1). This observation suggests that it is the differential expression of commonly detected proteins rather than the presence or absence of specific proteins that underlie site-specific plaque development. However, it also highlights a limitation of this microdissection approach, as the dissected plaques included the underlying medial and adventitial layers of vessel wall. Consequently, major structural proteins of the normal aortic wall (e.g. Myl6b and elastin (Eln)) that were detected within plaque samples likely originate, at least in part, from the underlying media.

One limitation of this study is the lack of immunohistochemical staining of specific proteins to validate the proteomics data. Although such analyses are difficult to combine with the current workflow, future studies should seek to validate key findings using complementary approaches. A further limitation is that this study was restricted to animals of a single age and sex, despite age and sex being important determinants of atherosclerotic plaque development. Application of this spatial aortic analysis workflow across multiple ages, sexes, and durations of WD feeding could provide important insight into the temporal sequence of the observed molecular changes and their sex-specific regulation. The identification of a molecular disease gradient across the aortic arch also raises the possibility of using this model to investigate therapeutic effects on different stages of atherosclerosis within a single animal. Although the present study lacks the resolution afforded by single cell or spatial proteomics approaches (e.g. [16, 51, 52]), it offers an attractive balance between spatial resolution, proteome depth, and experimental throughput. By enabling comprehensive proteomic profiling of numerous vascular regions across multiple animals, this workflow provides a practical framework for investigating the molecular characteristics of site-specific atherosclerotic plaque development and progression.

## Supporting information

Supplementary Table 1 Top 50 protein loadings of PC1

Supplementary Table 2 All proteins quantified

Supplementary Table 3 Enrichment analyses results

## Abbreviations

Apob: apolipoprotein B-100
ApoE^−/−^: apolipoprotein E-deficient
BH: Benjamini-Hochberg
Cd5l: CD5 antigen-like
Col8a1: collagen VIII alpha-1 chain
DIA-PASEF: data-independent acquisition with parallel accumulation-serial fragmentation
Eln: elastin
ECM: extracellular matrix
FDR: false discovery rate
Fn1: fibronectin
GSEA: gene set enrichment analysis
iBAQ: intensity based absolute quantification
Icam1: intercellular adhesion molecule 1
IGF: insulin-like growth factor
Itgb2: integrin beta-2
LC-MS/MS: liquid chromatography-tandem mass spectrometry
Lgals3: galectin-3
Lpl: lipoprotein lipase
Myl6b: myosin light chain 6B
Ngp: neutrophilic granule protein
NK cell: natural killer cell
NOS: nitric oxide synthase
PC1: principal component 1
PCA: principal component analysis
Pxdn: peroxidasin
S100a9: protein S100-A9
SPEED: Sample Preparation by Easy Extraction and Digestion
Vcam1: vascular cell adhesion molecule 1
WD: Western diet

## Data availability

All data are available within the article or Supplementary Data, except for the raw mass spectrometry data which have been deposited to the ProteomeXchange Consortium via the PRIDE partner repository [53] with the dataset identifier PXD080846. (Reviewer access token: KeZXYSsYXqzZ)

## Author Contributions

MJD, LFG and KVJ generated the hypothesis, designed and conceived the project. KVJ and LFG conducted experiments. KVJ, CC, MJD and LFG analysed, visualized and interpreted the data. KVJ, CC, MJD and LFG wrote the manuscript, provided intellectual input and edited the manuscript. All authors have given approval to the final version of the manuscript.

## Acknowledgements

We thank Ida-Mari Henriksen (Department of Clinical Biochemistry, Rigshospitalet, Copenhagen) for help with sacrificing the animals. The graphical abstract and Fig. 1B were created in BioRender under license.

