## Supplementary Data for "Location-dependent proteomics of the aorta reveal an atherosclerotic disease gradient shaped by hemodynamics"

* Corresponding authors

**Supplementary Data**

**Supplementary materials and methods**

**Sample preparation for proteomic analysis**

Proteins were extracted from the dissected aorta samples using the Sample Preparation by Easy Extraction and Digestion (SPEED) protocol [1]. 80% trifluoroacetic acid (TFA) was added to the samples and incubated for 30 min at room temperature, followed by incubation at 70°C for 10 min. Samples were then neutralized with 2 M Tris base using 8 times the volume of TFA used. Proteins were reduced and alkylated with 12 mM tris(2-carboxyethyl)phosphine and 47 mM 2-chloroacetamide for 10 min at 95°C, then digested with 1 µL of 0.05 µg LysC for 1 h at 37°C followed by adding 1 µL of 0.1 µg trypsin overnight at 37°C. All of the steps above were performed in a one-pot workflow in a PCR plate to prevent sample loss. Samples were acidified with 10% TFA (in water) and purified by solid-phase extraction using stage tips prepared with Affinisep AttractSPE C18 membranes [2]. The stage tips were conditioned with 100% methanol and washed with 0.1% TFA. Peptides were eluted with 50% acetonitrile (ACN) / 0.1% formic acid (FA) in water, dried down and stored at -80°C until analysis by LC-MS/MS.

**Liquid chromatography-tandem mass spectrometry (LC-MS/MS)**

Samples were reconstituted in 25 µL of 5% FA in water and analysed using a Dionex Ultimate RSLCnano chromatography system (Thermo Fisher) coupled to a timsTOF Pro (Bruker) mass spectrometer operated in a data-independent acquisition with parallel accumulation serial fragmentation (DIA-PASEF) mode [3]. 5 µL of sample was loaded on an Ion Opticks Aurora C18 column (15 cm x 75 µm, 1.7 µm particle size) at 35 °C with the peptides eluted using a solvent gradient of 0.1% FA in water (Solvent A) and 0.1% FA in ACN (Solvent B) over 22 min at a flow rate of 400 nL min^-1^. The gradient elution consisted of 4-25% B (0-14 min) and 25-85% B (14-14.5 min), followed by washing of the column with 85% B (14.5-18 min) and re-equilibration with 85-4% B (18-18.5 min) and 4% B (18.5-22min).

**Data analysis**

The DIA-PASEF data was searched against the *Mus musculus* UniProt reference proteome containing all reviewed proteins (UP000000589, downloaded 2024-12-12), including common contaminants, using DIA-NN version 1.9.2 in library-free mode [4]. Deep learning was used to create an *in silico* trypsin (Lys/Arg) digest of database with maximum 1 missed cleavage. Cysteine carbamidomethylation was enabled as a fixed modification together with protein N-terminal methionine excision, while methionine oxidation and N-terminal acetylation were set as variable modifications (maximum 1). Other settings used were; peptide length 7-35, precursor charge 2-4, precursor m/z 300-1000, fragment m/z 100-1700, false discovery rate (FDR) 1%, mass accuracy of 20/10 ppm (MS1/MS2), and peptidoforms, match-between-runs (MBR), and 'No shared spectra' enabled. The matrix of MaxLFQ protein group quantities was used for further analysis where contaminant proteins from other species as well as keratins were removed.

DIA-Analyst (version 0.10.3; https://analyst-suites.org/apps/dia-analyst) was used for standardised downstream statistical analysis of the protein-level data. Proteins with a high proportion of missing values were filtered out (protein retained if present in at least 3 out of 5 replicates in one group) and protein intensities were log_2_-transformed. For the differential expression analyses, protein-wise linear models combined with empirical Bayes statistics were used. The limma package from R Bioconductor was used to generate a list of differentially expressed proteins for each pair-wise comparison. Proteins were considered significantly regulated if the Benjamini-Hochberg (BH) [5] adjusted *p*-value was < 0.05 and log_2_ fold change > 1.

Further data analysis and visualisation was performed using R (version 4.5.2). Gene Set Enrichment Analysis (GSEA) was carried out using the clusterProfiler and msigdbr packages with the log_2_ fold changes from DIA-Analyst as input, and a BH adjusted *p*-value cut off < 0.05 [6]. The Molecular Signatures Database (MSigDB) [7-9] collections used for GSEA were the Reactome pathway [10] and cell type signature gene sets [11-13]. Intensity based absolute quantification (iBAQ) values were calculated as described previously [14]. In brief, for each protein, all matching proteotypic peptide intensities were summed and divided by the number of theoretically observable peptides from the *in silico* trypsin digest. The full DIANN report was used for this together with the QFeatures package. Matrisome annotation of the proteins was performed using the MatrisomeAnalyzeR package [15].
